# Thermal niches of specialized gut symbionts: the case of social bees

**DOI:** 10.1101/2020.06.16.155309

**Authors:** Tobin J. Hammer, Eli Le, Nancy A. Moran

## Abstract

Responses to climate change are particularly complicated in species that engage in symbioses, as the niche of one partner may be modified by that of the other. We explored thermal traits in gut symbionts of honeybees and bumblebees, which are vulnerable to rising temperatures. *In vitro* assays of symbiont strains isolated from 16 host species revealed variation in thermal niches. Strains from bumblebees tended to be less heat-tolerant than those from honeybees, possibly due to bumblebees maintaining cooler nests or inhabiting cooler climates. Overall however, bee symbionts grew at temperatures up to 44 °C and withstood temperatures up to 52 °C, at or above the upper thermal limits of their hosts. While heat-tolerant, most strains of the symbiont *Snodgrassella* grew relatively slowly below 35 °C, perhaps because of adaptation to the elevated body temperatures that bees maintain through thermoregulation. In a gnotobiotic bumblebee experiment, *Snodgrassella* was unable to consistently colonize bees reared below 35 °C under conditions that limit thermoregulation. Thus, host thermoregulatory behavior appears important in creating a warm microenvironment for symbiont establishment. Bee-microbiome-temperature interactions could affect host health and pollination services, and inform research on the thermal biology of other specialized gut symbionts, such as those of humans.

## Introduction

Earth’s climate is rapidly warming, and there is an urgent need to understand how organisms will respond [1, 2]. One factor complicating such predictions is the role of interspecific interactions [3, 4], and, in particular, symbiosis [5, 6]. Many organisms closely associate with one or more distantly related partners that have highly distinct physiologies, as in the case of animal or plant hosts and their microbiomes. Hosts and microbes are likely to have different responses to temperature; yet, if they are mutually dependent, the combined niche is restricted to that of the more sensitive partner. Furthermore, thermal niches can themselves evolve in response to symbiotic lifestyles. For example, obligate endosymbionts that undergo strong population bottlenecks during transmission may evolve unstable, easily denatured proteins as a consequence of mutation accumulation, leading to heat sensitivity [7–9]. Both of these factors may constrain the combined thermal niche of strongly symbiont-dependent organisms [6, 10, 11]. There is evidence for symbiont-imposed constraints on host thermotolerance in a variety of invertebrates such as aphids, ants, stinkbugs, corals, and sponges [12–16]. However, the wider prevalence of this phenomenon is unclear, and, in general, we do not know how microbiomes will influence host responses to climate warming.

The eusocial corbiculate bees (hereafter “social bees”) are a particularly important group in which to study symbiont thermal niches and their effects on hosts. This clade, comprising honey bees (*Apis*), bumblebees (*Bombus*) and stingless bees (Meliponini), is host to anciently associated, host-specialized, and beneficial gut microbiomes [17, 18]. Social bees are also key pollinators in both agricultural and natural ecosystems, but many are declining [19, 20]. For bumblebees in particular, rising temperatures have been identified as a driver of range shifts and population declines in some species [21, 22]. If gut symbionts are sensitive to heat stress, the microbiome could be one route through which climate change impacts bee health. Furthermore, because social bees as a whole exhibit extensive strain-level diversity in their microbiomes (e.g., [17, 23]), strain variability in thermotolerance could partially underlie corresponding variability among hosts, as was recently shown for endosymbionts of aphids [24].

Social bees present a uniquely complex thermal environment for their microbiome, making it challenging to predict their symbionts’ thermal traits. They are not strictly poikilothermic; rather, they facultatively regulate the temperature of both their bodies (and individual body parts) as well as their shared nests [25–27]. These microenvironments are partially buffered from external fluctuations in temperature, but to a degree that is highly dynamic among individuals, over time and space, and across the bee phylogeny. Even within a single nest, the microbiome is distributed across individuals that vary in behaviors such as foraging or brood incubation, which involve changes in host body temperature [28–30]. Furthermore, social bee species regulate their nest temperatures to different set-points and exhibit different overwintering strategies [27, 31, 32]. For example, the microbiomes of temperate-zone bumblebees must overwinter within diapausing queens, while the microbiome of *Apis mellifera* is transmitted by a cluster of active, heat-generating workers [33, 34].

As in gut symbionts generally, symbionts of social bees experience a brief *ex vivo* phase during transmission, potentially imposing selection on thermal traits. The gut microbiome is transmitted via a fecal-oral route, usually between nestmates within a hive [35, 36], but horizontal transmission between bee species has also been inferred [17, 34]. Although the symbionts cannot grow under ambient oxygen levels outside the bee gut [18], the ability to tolerate thermal stress while on flowers or other external habitats could influence horizontal transmission rates and thus patterns of biogeography and host specificity. All of these factors add up to a complex selective landscape—even within a single host species—involving different castes, seasons, and *ex vivo* phases.

Very little is currently known about the thermal biology of social bee microbiomes. Recent work on *Bombus impatiens* has shown that, once established in the gut, core symbionts are relatively robust to temperatures from 21–37 °C [37]. However, honeybees and bumblebees emerge largely symbiont-free as adults and must acquire their symbionts from nestmates [35, 36, 38]. It is not known if the colonization process, a crucial phase for both hosts and symbionts, is more temperature-sensitive than maintenance of an established symbiosis. Furthermore, there has been no comparative work investigating how symbiont thermal traits have evolved across social bees. Social bee taxa maintain different nest temperatures and use different strategies to overwinter and to establish new colonies. They also occupy a climatically diverse range of environments, from arctic and alpine habitats to tropical forests [39]. This variability may impose divergent selection on symbiont thermal traits, with potential feedbacks on the thermal tolerance of the hosts themselves.

We used common garden experiments (*in vitro*) to characterize symbiont thermal niches with a culture collection of *Snodgrassella* and *Gilliamella*, two bacterial species that are ubiquitous across honeybees and bumblebees [17]. Symbionts of these two bee lineages belong to deeply divergent clades and appear restricted to their native host [17, 40]. We measured the thermal limits to growth from 12-48 °C, the ability to tolerate a brief heat exposure up to 52 °C, and growth rates at 28 °C versus 35 °C. 28 °C is within the range of brood nest temperatures reported for some bumblebee species [41, 42] (though not all [43]), while honeybee brood nest temperatures are typically ~33–36 °C [31, 32, 44], at least for *Apis mellifera* and *A. cerana*.

We hypothesized that for all three metrics, honeybee-associated strains would exhibit higher thermotolerance than bumblebee-associated strains, as a result of adaptation to a generally warmer host (nests and bodies) and external environment. We also examined whether the thermal environment impacts *Snodgrassella* establishment in the bumblebee *Bombus impatiens*. This experiment examined whether the colonization process is particularly vulnerable to thermal stress; our initial hypothesis was that an abnormally high ambient temperature for *B. impatiens* (35 °C) would impair symbiont acquisition. Our findings provide a foundation for future work on the thermal ecology of bee gut microbiomes and raise new questions about the role of host thermoregulatory behavior in mediating symbiosis.

## Materials & Methods

### Culture collection

Strains of *Snodgrassella alvi* and *Gilliamella* spp. used in this study were collected as described in refs. [40, 45, 46]. To our knowledge, no strains had undergone significant passaging in the lab since the original isolates were obtained, with the exception of *Snodgrassella alvi* strain wkB2. Bumblebee host species were categorized into high-elevation or low-elevation species for Figs. S1, S2 following range descriptions in ref. [47].

### Thermal limits assay

From glycerol stocks, strains of *Snodgrassella* or *Gilliamella* were cultured on Columbia blood agar plates under 5% CO2 and 35 °C for 2 d. These were then restreaked, and the overnight cultures were resuspended in Insectagro medium (Corning) and adjusted to an OD_600_ of 0.5. We further diluted these cell suspensions 1/20 and spotted 10 μl in triplicate onto the surface of fresh Columbia plates. Negative controls (10 μl Insectagro spotted onto plates) were included to ensure there was no background contamination. Given that the same OD can correspond to different densities of viable cells, we also quantified the corresponding colony-forming unit (CFU) count of the inoculum for each strain. Plates were incubated at a range of temperatures under 5% CO2, from 12–48 °C in increments of 4 °C. After 48 h, we scored these plates for whether visible biomass was present or absent. If a strain could not grow at or below 28 °C, it was not tested at colder temperatures. For example, strain wkB339 could not grow at 28 °C so it was not further tested at 24 °C or below.

We used logistic regression (implemented in R [48] as a generalized linear model with binomial error and logit link function), to test whether host genus (*Apis* vs. *Bombus*) predicted the ability of symbionts to grow at 44 °C, the upper thermal limit across strains in this assay. To account for the potential influence of starting inoculum size on the probability of growth, we used the same approach, but with log-transformed CFU counts for each strain as a predictor.

### Heat stress assay

*Snodgrassella* cultures were initially prepared as above. We then resuspended overnight cultures in Insectagro, adjusted them to an OD_600_ of 0.5, and further diluted 1/20 in 200 μl in PCR plates. Cells were subjected to a 1 h heat stress treatment using a thermocycler with a temperature gradient from 35.4 °C to 51.6 °C (3 technical replicates per strain per temperature). They were then transferred to 96-well cell culture plates (Corning) in a microplate reader (Tecan) with 5% CO2 and 35 °C. Growth was monitored by OD_600_ readings taken every 3 h for 66 h. We included blank wells (Insectagro only) as negative controls and subtracted their OD_600_ values from those of the cultures.

### Thermal performance assay

*Snodgrassella* cultures were prepared as for the heat stress assay, except that they were transferred directly into 96-well cell culture plates in the microplate reader (without the thermocycler step) and incubated at either 28 °C or 35 °C for 72 h. We fit logistic curves to the data using the growthcurver package [49] and used a two-way ANOVA to test for effects of incubation temperature and host genus on the intrinsic growth rate (*r*). Linear regression was used to test whether starting inoculum size (log-transformed CFU counts) predicted growth rate.

### Colonization experiment

To obtain gnotobiotic bumblebees, we collected clumps of pupal cocoons from four separate commercial colonies of *Bombus impatiens* (Koppert USA). We then surface-sterilized the clumps in diluted bleach (0.2% NaOCl) for 90 s as described previously [50, 51] to minimize contamination of the emerging workers. We maintained the sterilized cocoon clumps in sterile conditions in a growth chamber at 35 °C and monitored them daily for adult emergence. Newly emerging adults were transferred to sterile vials and randomly assigned to *Snodgrassella* or buffer-only treatments. The former were fed with 10 μl of filter-sterilized sugar syrup (50% v/v) containing ~10^6^ cells of *Snodgrassella alvi* strain wkB12, following [40]. We prepared new *Snodgrassella* inocula daily from overnight cultures; these were not continuously propagated but rather independently obtained from the same frozen stock of wkB12. Negative-control bees were fed an identical solution but without cells. All bees were monitored to ensure that they consumed the entire 10 μl of inoculum.

We then transferred bees to sterilized 16 oz plastic containers (Dart Container Corp.) in groups of 2-3 as microcolonies [52]. Bees within a microcolony were assigned to the same treatment and were obtained from the same source colony. Each container was provided with 10 ml of sterile 50% sugar syrup and 500 mg of sterile pollen dough (ground gamma-irradiated honeybee pollen mixed with sterile 50% sugar syrup). Microcolonies were reared in incubators at 29 °C, representing a typical bumblebee rearing temperature [37, 52, 53], or 35 °C, a temperature typical of *Apis mellifera* hives [32]. The pollen lump was replaced on the third day of rearing.

After 5 d, bees were briefly anaesthetized in ice and used for gut dissections. The gut (including hindgut and midgut) was removed from each bee, homogenized with a sterile plastic pestle, and resuspended in 1 ml of Insectagro. This homogenate was serially diluted and plated on Columbia blood agar. We counted CFUs after 2 d of incubation at 35 °C in 5% CO2. In performing CFU counts of the focal *Snodgrassella* strain, we noticed the occasional presence (in 15% of bees overall) of an unidentified bacterium. This contaminant had distinct colony morphology and exhibited slow growth and hemolysis. There was no significant association between the inoculum or the rearing temperature with the presence of the contaminant (logistic regression; inoculum *p* = 0.19, temperature *p* = 0.066). This may have been a core bee gut symbiont not completely removed from the cocoon surface by the sterilization treatment.

We used a linear mixed-effects model with the nlme package [54], treating colony source as a random effect, to test whether rearing temperature was associated with log-transformed *Snodgrassella* CFU counts. We also used logistic regression (as described above) to test whether bee survival was predicted by the inoculum treatment or by rearing temperature.

## Results

We first used an *in vitro* assay to measure the temperature limits of growth of a collection of *Snodgrassella* and *Gilliamella* strains isolated from honeybees and bumblebees. While all strains could grow at 40 °C (Fig. S1) and none at 48 °C, they varied in their ability to grow at 44 °C (Fig. 1). Bumblebee strains were significantly less likely to be able to grow at this temperature than honeybee strains (logistic regression, *p* = 0.015).

**Figure 1.**
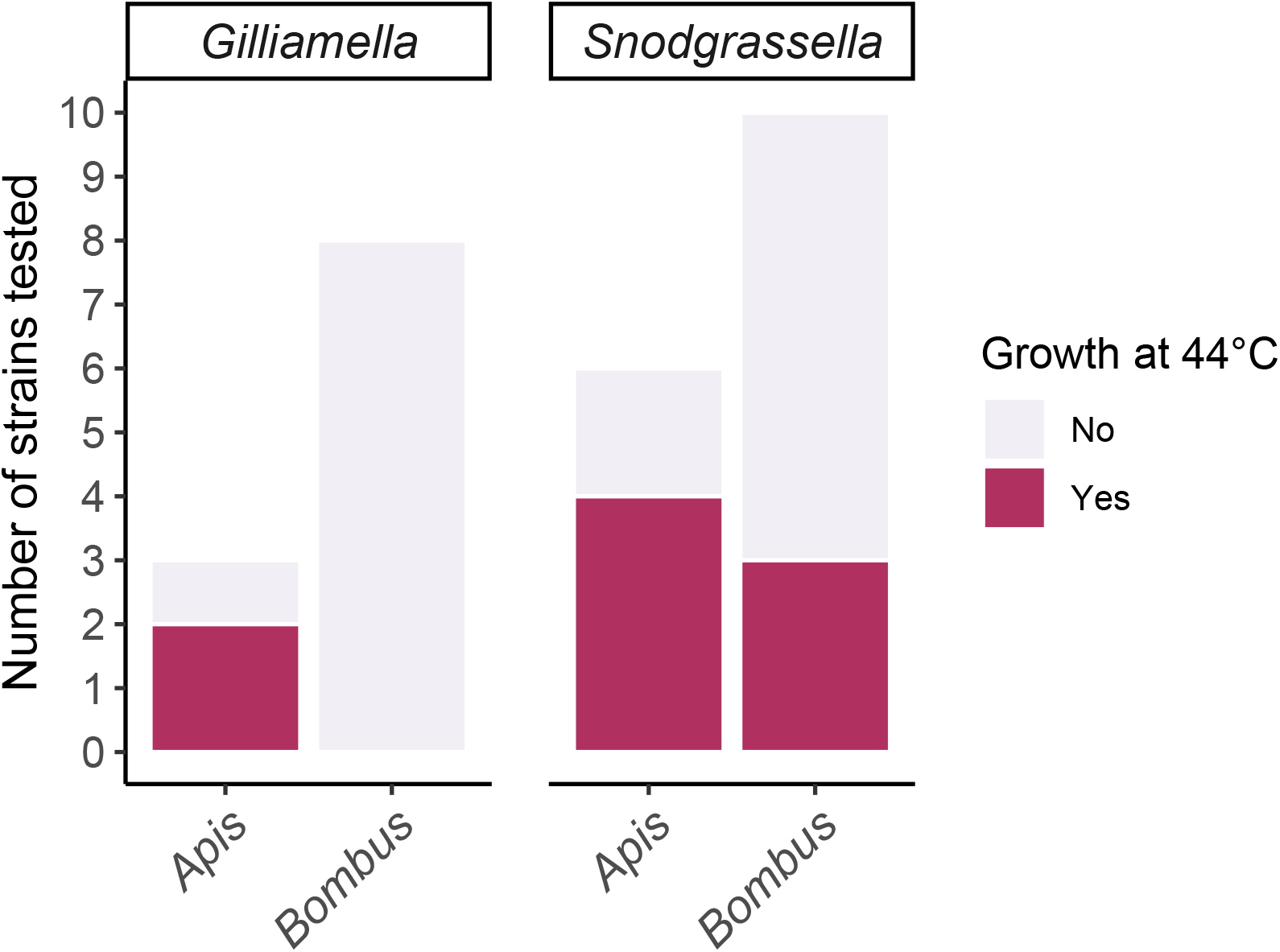
Ability of strains of two core bee gut symbionts, *Gilliamella* and *Snodgrassella*, to grow after 48 h incubation on solid media at 44 °C. Honeybee (*Apis*) strains tend to be better able to grow at this temperature than bumblebee (*Bombus*) strains. Thermal limits broken down by host species of origin are shown in Fig. S1.

There was also substantial strain-level variation in lower thermal limits, especially for *Snodgrassella* (Fig. S1). However, probability of growth at the lowest temperature tested (12 °C) was positively associated with starting inoculum size (logistic regression, *p* = 0.0047), which was not the case for growth at 44 °C (logistic regression, *p* = 0.74). Hence, we regard the lower limit data as minimum estimates; many of these strains are likely to be able to grow at lower temperatures than indicated in Fig. S1, especially with larger starting inocula or a longer assay duration.

In addition to characterizing the heat-sensitivity of symbiont growth under a constant temperature, we also sought to determine the ability of bee gut symbionts to tolerate short-term exposure to more extreme heat. Exposure to temperatures above 35 °C for 1 h clearly delayed subsequent growth *in vitro* (Fig. 2). Overall, however, *Snodgrassella* appears to be quite heat-tolerant in that all six strains assayed could recover from exposure to 48.7 °C. The exact limit was variable among strains. Although larger strain sample sizes are needed for a conclusive comparison, the three *Apis-*associated strains tended to be slightly more heat-tolerant than the three *Bombus*-associated strains (Fig. 2). Strain robustness to short-term exposures was not simply a function of the initial inoculum size. For example, the only two strains that could recover following exposure to 51.6 °C, wkB9 and wkB237, had the lowest inoculum sizes.

**Figure 2.**
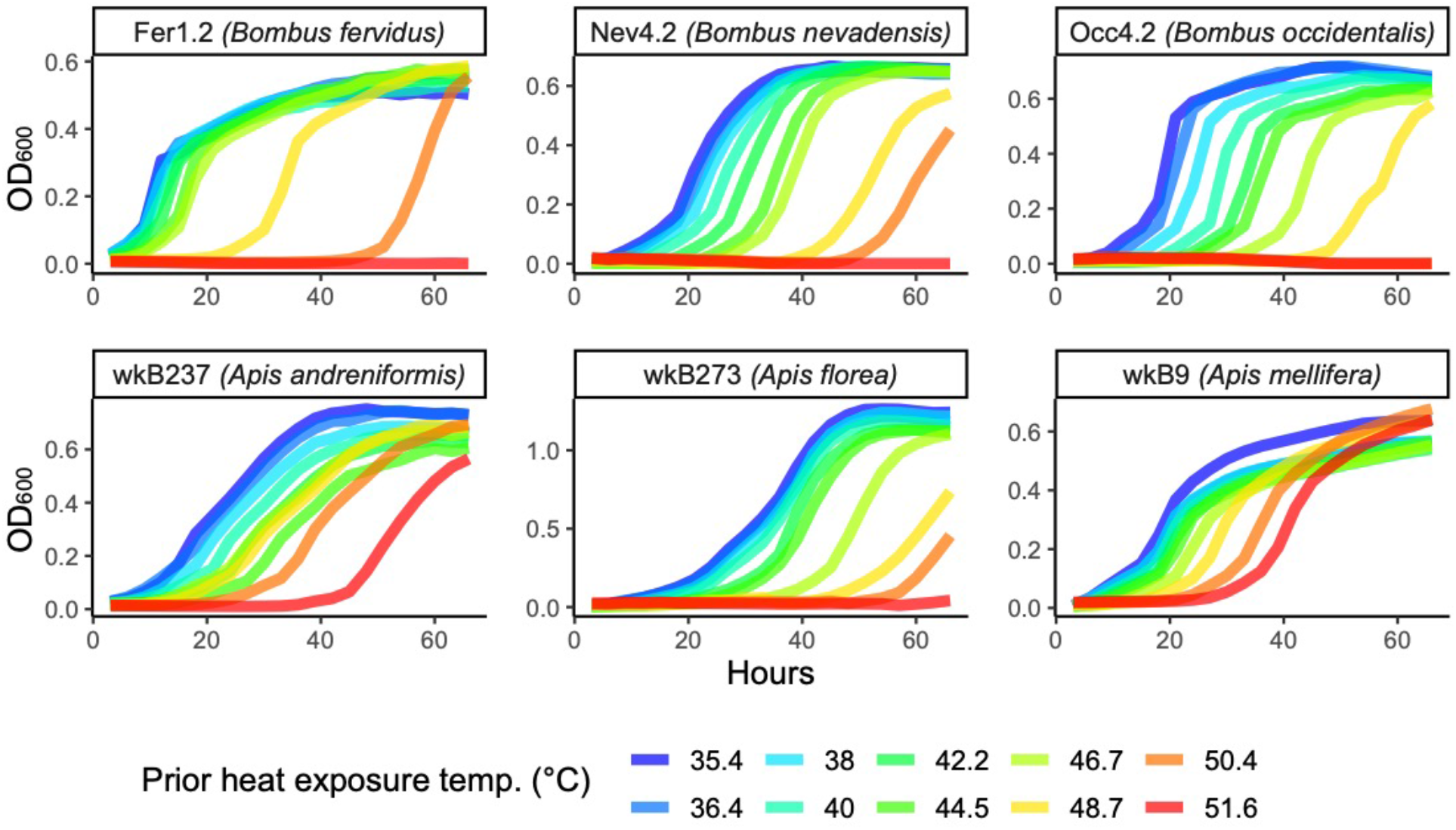
*In vitro* growth of the core bee gut symbiont *Snodgrassella* following short-term exposure to high temperatures. Curves represent growth at 35 °C and 5% CO2 following a 1 h heat stress treatment applied using a gradient thermocycler, showing the mean OD_600_ values of three replicates per strain per temperature. The host species from which each strain was isolated is indicated. *Snodgrassella* is generally robust to high temperatures, though tolerance varied among strains. Only two *Apis*-associated isolates were able to recover from 51.6 °C, the maximum temperature tested.

Even when thermal limits do not vary among symbiont strains, the thermal optima could vary. To address this, we conducted *Snodgrassella* growth assays in liquid media at 28 °C and 35 °C, representing temperatures closer to typical *Bombus* or *Apis* nest temperatures, respectively. Inoculum size (number of CFUs at the start of the growth assay) did not predict growth rate at either 28 °C or 35 °C (linear regression, *p* = 0.69 and 0.82, respectively). Interestingly, bumblebee-associated *Snodgrassella* typically had higher growth rates than honeybee-associated *Snodgrassella* (Fig. 3; two-way ANOVA, *p* = 0.0085). We found no significant interaction between host genus and temperature (two-way ANOVA, *p* = 0.24), as would be expected if symbiont growth were differentially adapted to the temperatures closest to those of their hosts’ nests. Instead, growth rates of most strains were higher at 35 °C (Fig. 3; two-way ANOVA, *p* = 0.0059). Among bumblebee strains, those from host species inhabiting higher elevations were not conspicuously better able to grow at the cooler temperature, 28 °C, than those from low-elevation hosts (Fig. S2). In fact, the only strains that exhibited higher growth rates at 28 °C than at 35 °C were derived from *Apis mellifera* (Fig. S2).

**Figure 3.**
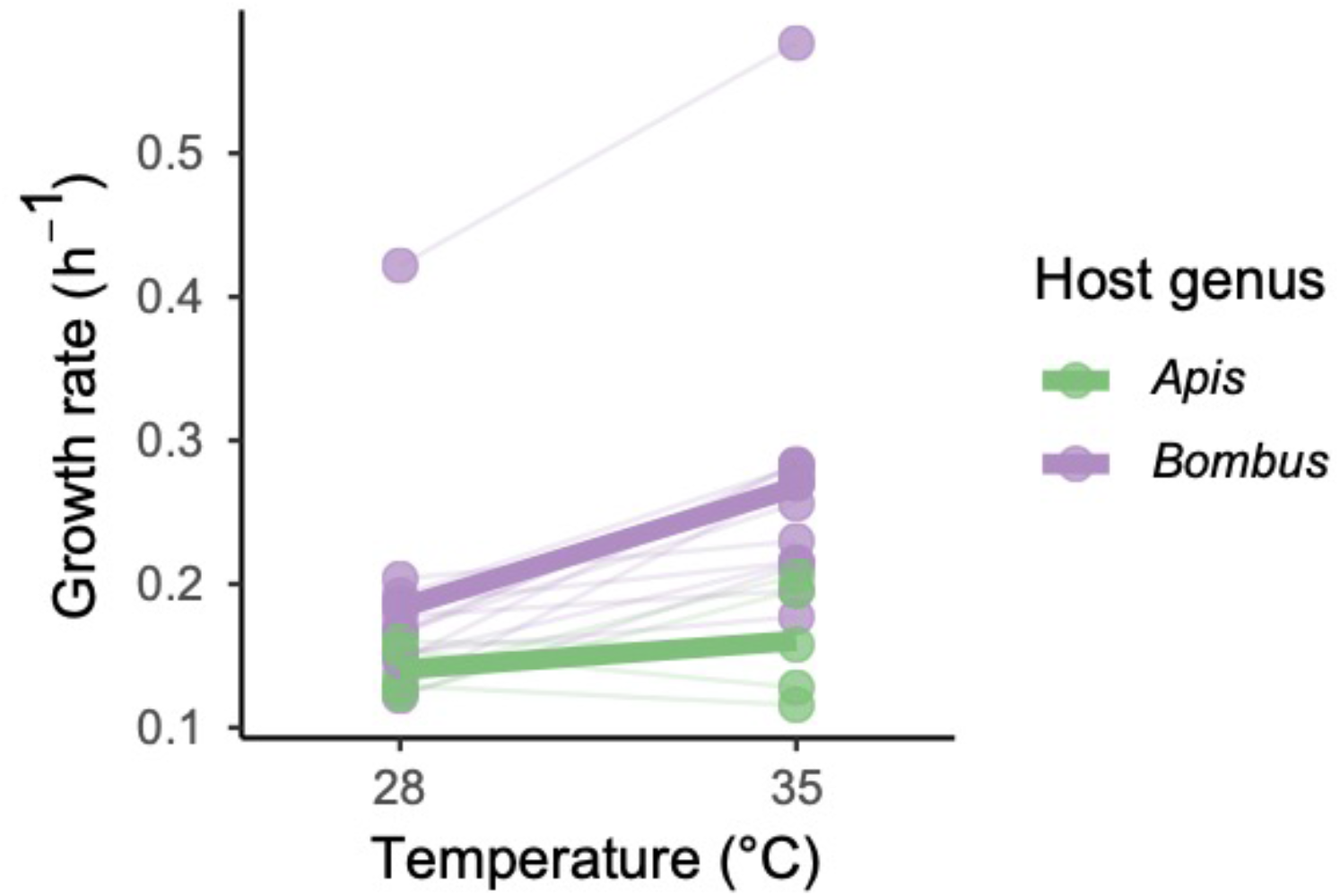
*In vitro* growth rates of *Snodgrassella* strains at 28 °C and 35 °C. Dots connected by thin lines represent the maximum growth rates of each strain, colored by whether the strains were isolated from honeybees (*Apis*) or bumblebees (*Bombus*). Thick lines connect the median growth rate for *Apis* or *Bombus* strains at each incubation temperature. All bumblebee *Snodgrassella* grow faster at 35 °C, a temperature significantly higher than typical bumblebee nests. N = 5 *Apis* strains, N = 15 *Bombus* strains.

We next examined whether symbiont thermotolerance might influence fitness *in vivo* using a colonization experiment with *Snodgrassella alvi* strain wkB12 and gnotobiotic bumblebees. *In vitro*, this strain grows much more quickly at 35 °C as compared with 28 °C (Fig. 4B). No *Snodgrassella* colony-forming units (CFUs) were detected in any of the bees inoculated with sterile buffer (N = 15 reared at 29 °C, N = 16 reared at 35 °C). Among bees inoculated with a standardized dose of ~10^6^ *Snodgrassella* cells, gut colonization after 5 days was highly dependent on the thermal environment (Fig. 4A). Accounting for the different source colonies from which bees were obtained, CFU counts were significantly higher (by over 50-fold, comparing medians) in the 35 °C rearing treatment (linear mixed-effects model,*p* < 0.0001). Several bees reared at 29 °C had no detectable *Snodgrassella* cells.

**Figure 4.**
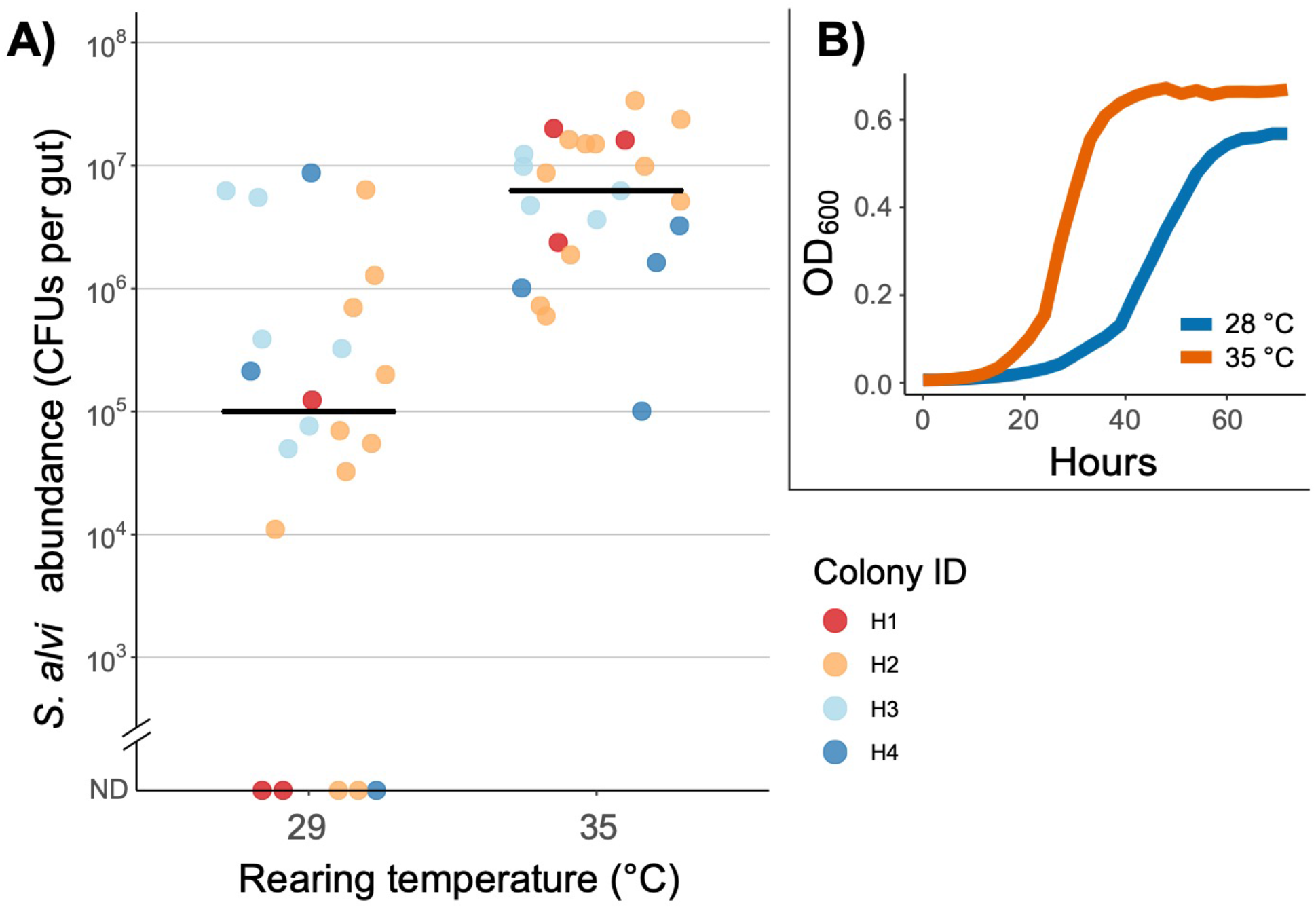
A) Effects of the thermal environment on colonization of *Snodgrassella alvi* wkB12 in gnotobiotic bumblebee workers (*Bombus impatiens*). Inoculated bees were maintained for 5 d at 29 °C, a typical temperature in *B. impatiens* nests (N = 22), or 35 °C (N = 23). Colors indicate the four replicate colonies that were used for the experiment, and black bars indicate the median *Snodgrassella* titer for each temperature treatment. Bees fed a sterile buffer and maintained under the same conditions had no detectable *Snodgrassella* colonization (not shown). CFUs = colony-forming units; ND = not detected. B) Growth of *Snodgrassella alvi* wkB12 *in vitro* when incubated at 28 °C versus 35 °C. Growth curves represent the mean OD_600_ values of three replicates per temperature.

We also examined whether bee survival was influenced by the experimental treatments. Overall, 92% of bees survived from adult emergence to dissection on day 5. The probability of survival was not affected by the inoculum (*Snodgrassella* vs. buffer alone) or the rearing temperature (logistic regression; inoculum *p* = 0.49, temperature *p* = 0.33) (Fig. S3). As only bees that survived for the entire 5-day rearing period were used for CFU counts, we were not able to examine whether gut colonization was predictive of survival.

## Discussion

We first used *in vitro* assays to characterize bee symbiont thermal niches in a common environment and without potential host-mediated effects. The experiments were based on a collection of isolates of *Snodgrassella* and *Gilliamella*, which are ubiquitous, keystone members of the social bee gut microbiome [17, 18]. Overall, we found that these symbionts are quite heat-tolerant relative to their hosts. All strains of both symbiont species can grow at a constant temperature of 40 °C, and many can grow at 44 °C (Fig. 1). Likewise, another core symbiont, *Lactobacillus bombicola*, has an optimal growth temperature around 40 °C [55]. In contrast, honeybees and bumblebees do not normally allow their nests to reach temperatures above 35 °C, which would harm brood development [31, 32, 41–43, 56].

Bees can, however, maintain higher body temperatures for brief periods while foraging. For example, the abdomen (which contains the gut microbes [18]) of bumblebees foraging in full sunlight may reach close to 40 °C [28]. The abdomen is also used to dissipate excess heat generated in the thorax [57]. We used short-term heat treatments of *Snodgrassella* to mimic this kind of temporary exposure, and found that bumblebee-associated strains could recover from an hour-long exposure to at least 48.7 °C, while honeybee-associated strains could recover from at least 50.4 °C (Fig. 2). In contrast, measured lethal limits (broadly defined) of bumblebees and honeybees are similar or lower, around ~40–46 °C [58, 59] and ~50 °C [60–62], respectively.

Bumblebees have experienced population declines partly linked to climate warming [21, 22]; in this study, we asked whether heat-sensitive symbionts could constitute one underlying mechanism. The comparative robustness of *Snodgrassella, Gilliamella*, and *Lactobacillus bombicola* to high temperatures *in vitro* suggests that the gut microbiome does not constrain bee tolerance of heat. Rather, other factors rooted in host physiology, behavior, and ecology likely explain the observed impacts of climate change on bumblebee populations [63–65]. However, further experiments *in vivo* are needed to conclusively test whether the microbiome plays a role in mediating effects of heat stress on bee populations.

*Snodgrassella* and *Gilliamella* are heat-tolerant not only compared to their hosts, but also compared to many insect endosymbionts. For example, aphids, weevils, carpenter ants, and stinkbugs all have obligate associations with highly heat-sensitive endosymbionts [10, 13, 14, 24, 66]. One trait these symbionts share is strict maternal inheritance; this transmission mode enforces a clonal population structure that results in genome degeneration and, ultimately, impaired heat tolerance [7–9]. In contrast, social transmission of gut symbionts permits the strain mixing and recombination that is more typical for free-living bacteria. While mostly vertically transmitted between colonies, bee gut symbionts likely maintain larger population sizes and undergo recombination more frequently than endosymbionts. All bee gut symbiont species are culturable outside the host [67] and possess genomes that do not exhibit the hallmarks of degenerative evolution [40].

Like many gut microbes, bee gut symbionts experience a brief but potentially important *ex vivo* phase during transmission. Selection for persistence on nest substrates or in the environment (e.g., flowers) may influence their heat tolerance. *In vivo* selective pressures related to the unique thermoregulatory behavior of social bees may further explain the broad thermal range of bee symbionts. A bee-inhabiting microbial population will experience a wide range of body temperatures as its hosts forage in the environment, thermoregulate nests, and overwinter. In the future, comparisons to non-bee-associated bacterial relatives (e.g., [68]) would be useful to reconstruct how evolution in social bees specifically has shaped the thermal niches of the bee gut microbiome.

While generally heat-tolerant relative to their hosts and to other bacterial symbionts of insects, *Snodgrassella* and *Gilliamella* strains do vary in thermal traits. This variability manifests as higher heat tolerance for honeybee versus bumblebee strains (Fig. 1, Fig. 2), a pattern that roughly matches the corresponding thermal traits of their hosts. For example, honeybees typically occupy tropical and subtropical environments (with the exception of introduced *Apis mellifera*), maintain warmer nests, have higher upper thermal limits, and do not undergo diapause in winter [26, 62, 69]. Our findings are consistent with previously observed correlations between symbiont thermotolerance and the local thermal environment [24, 70–72]. In the case of bees, even if divergent thermal niches of symbionts do not affect hosts, they could affect the potential for strains to successfully disperse between host colonies or even species, ultimately influencing their biogeography and degree of host specialization.

Surprisingly, we did not find such host-symbiont matching for *Snodgrassella* thermal performance (i.e., relative growth rate at two temperatures). Bumblebee strains uniformly grew faster at 35 °C, while most *Apis mellifera* strains grew slightly faster at 28 °C (Fig. 3, Fig. S2). This pattern is the opposite of what would be expected if the ambient temperature in active colonies primarily determines the optimal growth temperature of symbionts, because honeybees generally maintain warmer nests. We speculate that overwintering biology may explain the *A. mellifera*-derived strains’ comparatively higher growth rates at the cool assay temperature. Unlike bumblebees, in winter, *A. mellifera* workers form active clusters that maintain above-ambient, but cool temperatures (average ~21 °C; [32]).

What explains our observation that bumblebee-associated *Snodgrassella*, like *Lactobacillus bombicola* [55], grow faster at a temperature exceeding that of their nests (Fig. 3)? We suggest that the answer lies in the bee abdomen—the microenvironment inhabited by the symbionts, and a structure whose temperature is affected by host thermoregulatory behavior. Specifically, both workers and queens (and occasionally males [73]) of bumblebees incubate larvae and pupae by placing their abdomen directly onto brood structures and elevating its temperature to ~35 °C or above, exceeding ambient temperatures in the nest [74, 75]. Given that bumblebees spend much of their time performing this behavior [41, 74, 76]—especially foundress queens, the sole source of microbes for the colony—symbiont growth may be adapted to the locally heated conditions within the abdomen.

To further explore this possibility, we tested whether the warm-shifted growth preference of bumblebee-associated *Snodgrassella* (Fig. 3) might have fitness effects *in vivo*. We conducted an experiment on gnotobiotic *Bombus impatiens*, using conditions (microcolonies lacking brood) in which incubation behavior is limited, and thus abdominal temperatures are expected to more closely match the chosen ambient rearing temperatures (29 °C and 35 °C). Previous literature hinted that these temperatures might affect *Snodgrassella* colonization of *B. impatiens*. In one study, experimental inoculation of *Bombus impatiens* with *Snodgrassella* resulted in 100% colonization and consistently high titers [40], whereas a later study of *B. impatiens* reported erratic colonization and frequently low titers [51]. These studies differed in the temperature at which bees were reared, with the high colonization rate observed at 34 °C and erratic colonization at ~26 °C. In line with these previous studies, when we directly examined the effect of the thermal environment on *Snodgrassella* colonization, we found that colonization was variable and occasionally unsuccessful in bees reared at 29 °C (Fig. 4A).

One possible explanation for this result has to do with the fact that the focal strain used, *Snodgrassella alvi* wkB12, was not isolated from *B. impatiens* but rather *B. bimaculatus*. If bumblebees control colonization by other species’ or even colonies’ symbiont strains, as has been suggested [51], robust colonization at 35 °C may actually reflect a weakening of strain-specific filtering mechanisms at this unnaturally high rearing temperature. However, we currently lack evidence for such mechanisms.

Another explanation for this result is based in the thermal niche of bumblebee-associated *Snodgrassella* strains. *Snodgrassella alvi* wkB12, like the others tested, grows more quickly at 35 °C than 28 °C *in vitro* (Fig. 4B). In bees reared in microcolonies at 29 °C—which we expect to have similar abdominal temperatures because of restricted thermoregulatory behavior— ingested *Snodgrassella* cells may simply be unable to replicate quickly enough to establish a stable population before being lost to defecation. However, once established, they appear to be quite robust to a wide range of temperatures [37]. A subset of bees reared at 29 °C did acquire high *Snodgrassella* titers that exceeded the number of cells in the inoculum, implying replication in the gut (Fig. 4A). One possibility is that these individuals had begun to incubate the provided pollen lump, a behavior which has been observed in microcolonies (e.g., [77]). As a consequence, they may have maintained higher abdominal temperatures conducive to *Snodgrassella* colonization. Analogously, in a normal bumblebee nest, individual-level thermoregulatory behaviors may be important in enabling establishment of the gut microbiome.

We note that our *in vitro* and *in vivo* experiments were conducted on single isolates, which enables controlled assays of strain-level thermotolerance but may miss important community interactions that occur within the bee gut. Microbial thermotolerance can be modified by co-occurring microbes or viruses (e.g., [78–81]), and phage have recently been documented from the bee gut microbiome [82, 83]. Community interactions may also help explain why monocolonizations of *Snodgrassella* into bees reared at 29 °C or below have been erratic (Fig. 4A and [51]), while fecal transplants at these temperatures have not [36, 50]. Another difference between *Snodgrassella* colonization from cultured isolates and from fecal transplants is in the quantity and physiological state of the cells that bees ingest. These factors could also help symbionts overcome a temperature barrier to gut colonization.

## Conclusions

We have argued that the gut microbiome is probably not a major constraint on how social bees respond to heat stress. However, as their growth is somewhat cold-sensitive, symbionts could be affected by other kinds of environmental stressors that disrupt host thermoregulatory behavior. For example, neonicotinoid pesticides interfere with *B. impatiens* thermoregulation [84]. Pesticides could thus indirectly affect colonization by *Snodgrassella* and perhaps other key symbionts, a process we have shown to be temperature-sensitive. Whether symbionts indeed constrain (or even improve [85]) bee responses to heat and other stressors remains an important priority for future research, as bees and the pollination services they provide continue to face serious challenges.

Our findings are also relevant beyond bees. We have identified a potential feedback between host behavior and the microbiome, a topic that has recently garnered substantial interest [86, 87]. Specifically, we suggest that social bee thermoregulatory behaviors have provided elevated and buffered microenvironments (nests and individual bodies), shaping the evolution of their symbionts’ thermal niches. Additionally, there are parallels between bees and mammals, which also create warm gut environments and harbor socially transmitted gut symbionts. Bees could be a useful model to understand how these factors influence symbiont thermotolerance, a little-studied trait in humans and other mammals.

## Data Availability

Primary data and R code used for analyses and visualizations are publicly available at the figshare repository: https://doi.org/10.6084/m9.figshare.12486395.

## Acknowledgments

This research was supported by a postdoctoral fellowship (2018-08156) from the USDA National Institute of Food and Agriculture to TJH, a University of Texas Undergraduate Research Fellowship to EL, and National Institutes of Health award (R35GM131738) to NAM. We also thank S. Leonard and E. Powell for helpful advice and K. Hammond for assistance.

## Supplemental Figures

**Figure S1.**
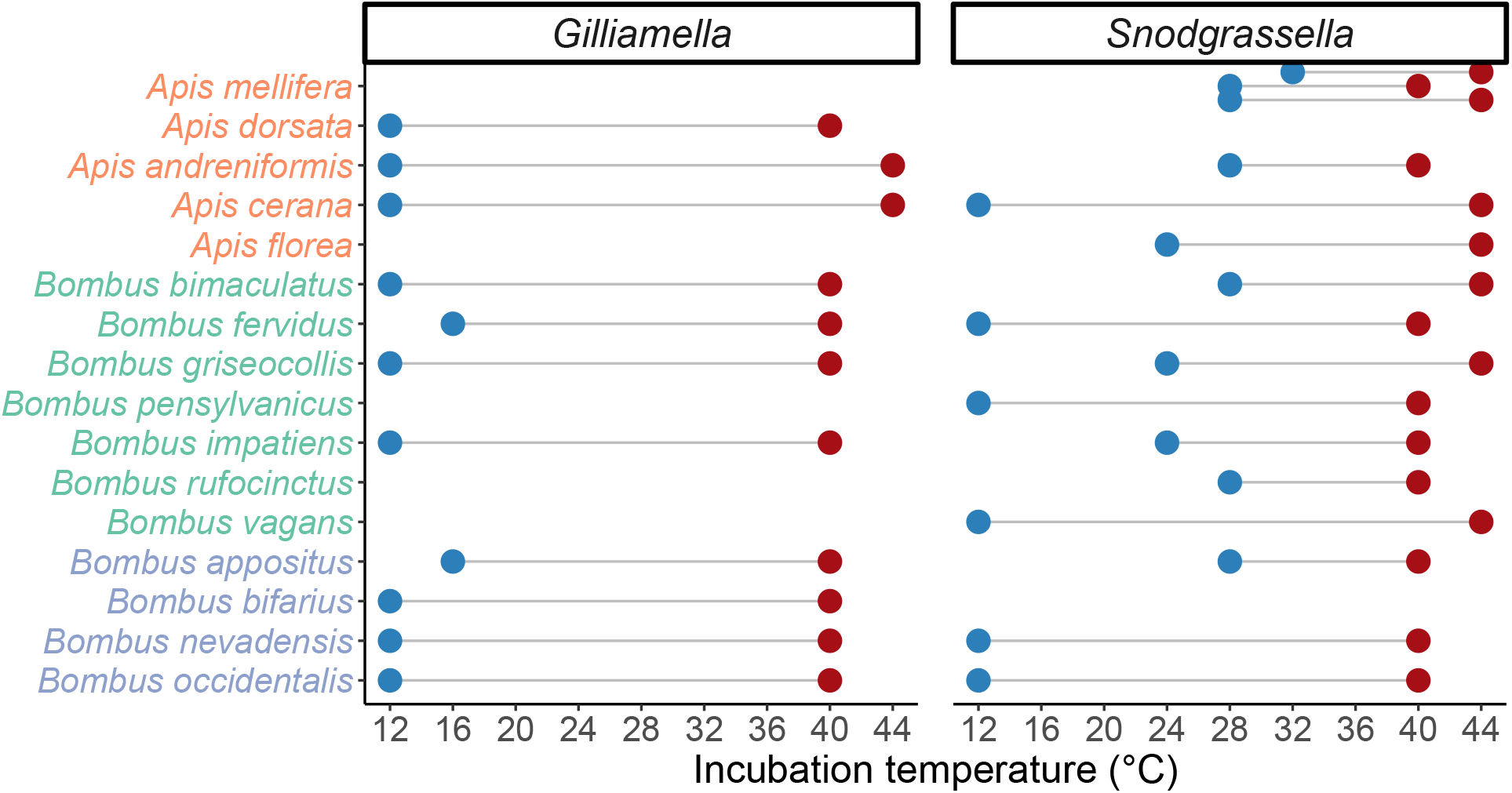
Thermal limits to growth *in vitro* for two bee gut symbiont species. Dots represent the minimum and maximum temperatures at which a given strain exhibited growth after 48 h of incubation on solid media. No growth was evident for any strain at 48 °C, and temperatures below 12 °C were not tested. Aside from *Apis mellifera*, from which three *Snodgrassella* isolates were assayed, one symbiont isolate per host species was assayed; note that blank spaces indicate untested host/symbiont combinations. The four bumble bee host species that typically inhabit high-elevation habitats are labeled in blue at the bottom of the plot. Lower limits to growth should be considered as minimum estimates of cold tolerance, given the sensitivity of this metric to the starting inoculum size (see Results).

**Figure S2.**
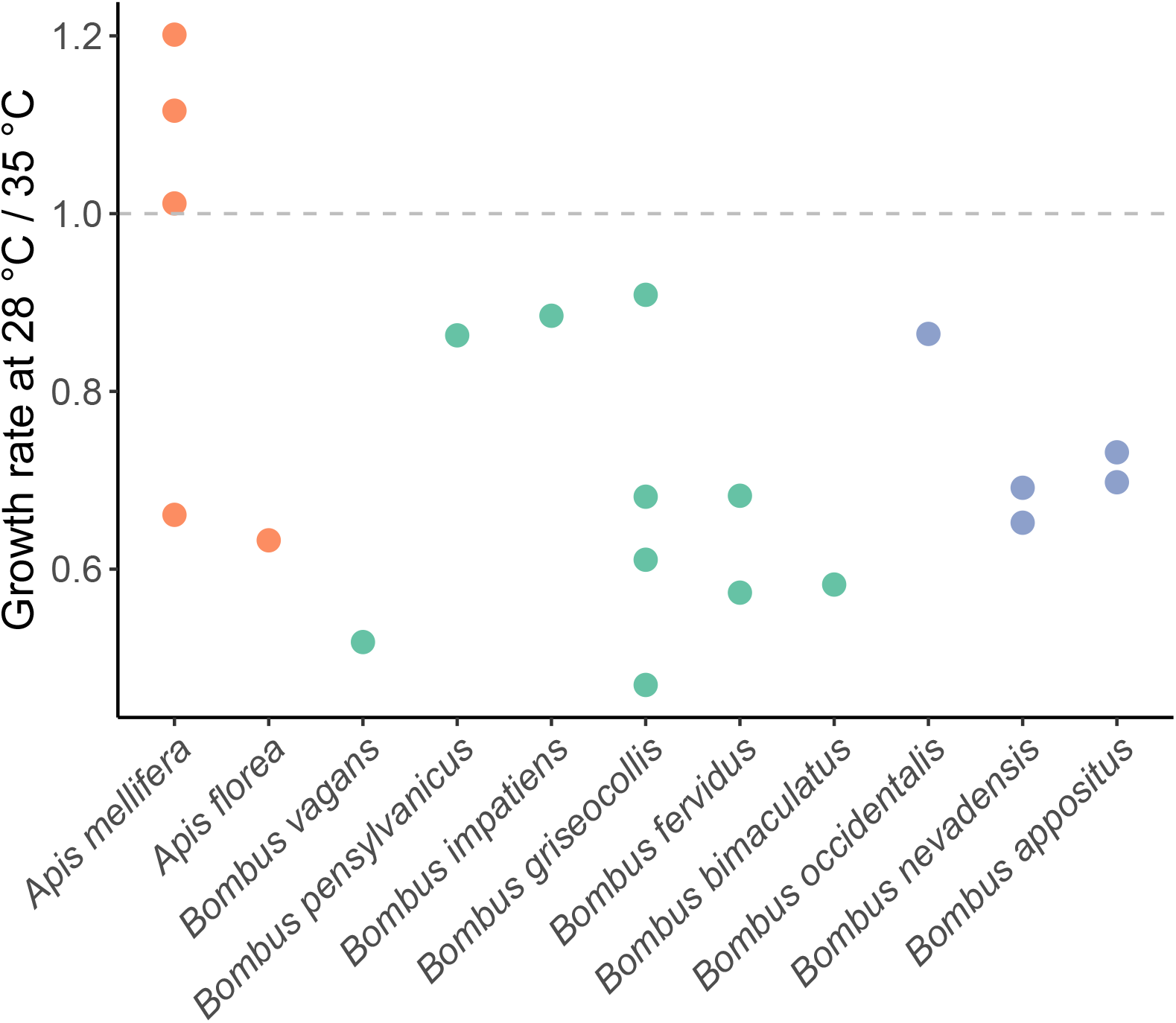
Growth rates of *Snodgrassella* at 28 °C versus 35 °C *in vitro*. The dashed line indicates equivalent growth rates between the two temperatures. Dots represent individual *Snodgrassella* strains, and strains from bumble bee species that typically inhabit high-elevation habitats are labeled in blue. The one *A. mellifera-derived* strain that grew faster at 35 °C is wkB2, which has been frequently cultured in the laboratory at 35 °C (see Methods). Its thermal performance may therefore reflect laboratory evolution.

**Figure S3.**
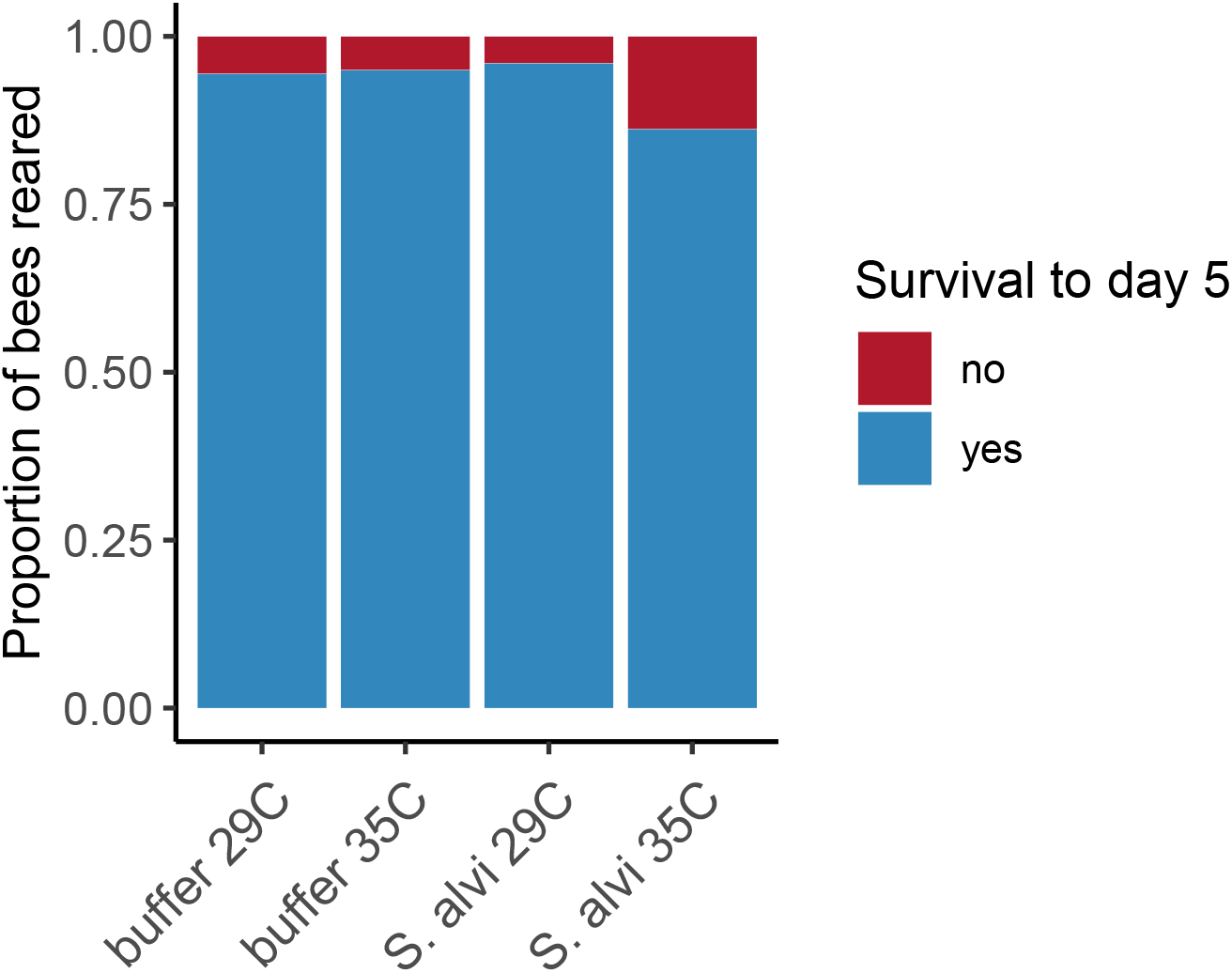
Survival outcomes of gnotobiotic bumble bees from adult emergence through five days of rearing. Number of bees inoculated with buffer alone: 18 at 29 °C, 20 at 35 °C. Number of bees inoculated with *Snodgrassella alvi* wkB12: 25 at 29 °C, 29 at 35 °C.

## References

1. Cahill AE et al. 2013 How does climate change cause extinction? Proc. R. Soc. B 280, 20121890. (doi:10.1098/rspb.2012.1890)

2. Urban MC. 2015 Accelerating extinction risk from climate change. Science 348, 571–573. (doi:10.1126/science.aaa4984)

3. Harrington R, Woiwod I, Sparks T. 1999 Climate change and trophic interactions. Trends in Ecology & Evolution 14, 146–150. (doi:10.1016/S0169-5347(99)01604-3)

4. Colwell RK, Dunn RR, Harris NC. 2012 Coextinction and Persistence of Dependent Species in a Changing World. Annual Review of Ecology, Evolution, and Systematics 43, 183–203. (doi:10.1146/annurev-ecolsys-110411-160304)

5. Kiers TE, Palmer TM, Ives AR, Bruno JF, Bronstein JL. 2010 Mutualisms in a changing world: an evolutionary perspective. Ecology Letters 13, 1459–1474. (doi:10.1111/j.1461-0248.2010.01538.x)

6. Wernegreen JJ. 2012 Mutualism meltdown in insects: bacteria constrain thermal adaptation. Current Opinion in Microbiology 15, 255–262. (doi:10.1016/j.mib.2012.02.001)

7. Melnikov SV, van den Elzen A, Stevens DL, Thoreen CC, Söll D. 2018 Loss of protein synthesis quality control in host-restricted organisms. Proc Natl Acad Sci USA 115, E11505–E11512. (doi:10.1073/pnas.1815992115)

8. Fares MA, Ruiz-González MX, Moya A, Elena SF, Barrio E. 2002 GroEL buffers against deleterious mutations. Nature 417, 398–398. (doi:10.1038/417398a)

9. Moran NA, McCutcheon JP, Nakabachi A. 2008 Genomics and Evolution of Heritable Bacterial Symbionts. Annual Review of Genetics 42, 165–190. (doi:10.1146/annurev.genet.41.110306.130119)

10. Corbin C, Heyworth ER, Ferrari J, Hurst GDD. 2017 Heritable symbionts in a world of varying temperature. Heredity 118, 10–20. (doi:10.1038/hdy.2016.71)

11. Moran NA. 2016 When obligate partners melt down. mBio 7, 7–9. (doi:10.1128/mBio.01904-16)

12. Dunbar HE, Wilson ACC, Ferguson NR, Moran NA. 2007 Aphid thermal tolerance is governed by a point mutation in bacterial symbionts. PLoS Biology 5, e96. (doi:10.1371/journal.pbio.0050096)

13. Fan Y, Wernegreen JJ. 2013 Can’t Take the Heat: High Temperature Depletes Bacterial Endosymbionts of Ants. Microb Ecol 66, 727–733. (doi:10.1007/s00248-013-0264-6)

14. Kikuchi Y, Tada A, Musolin DL, Hari N, Hosokawa T, Fujisaki K. 2016 Collapse of Insect Gut Symbiosis under Simulated Climate Change. 7, 1–8. (doi:10.1128/mBio.01578-16.Editor)

15. Weis VM. 2008 Cellular mechanisms of Cnidarian bleaching: stress causes the collapse of symbiosis. Journal of Experimental Biology 211, 3059–3066. (doi:10.1242/jeb.009597)

16. Webster NS, Cobb RE, Negri AP. 2008 Temperature thresholds for bacterial symbiosis with a sponge. ISME J 2, 830–842. (doi:10.1038/ismej.2008.42)

17. Kwong W et al. 2017 Dynamic microbiome evolution in social bees. Science Advances 3, e1600513. (doi:10.1126/sciadv.1600513)

18. Kwong WK, Moran NA. 2016 Gut microbial communities of social bees. Nature Reviews Microbiology 14, 374–384. (doi:10.1038/nrmicro.2016.43)

19. Cameron SA, Sadd BM. 2020 Global Trends in Bumble Bee Health. Annu. Rev. Entomol. 65, 209–232. (doi:10.1146/annurev-ento-011118-111847)

20. Potts SG. 2015 Global pollinator declines: Trends, impacts and drivers. Trends in Ecology and Evolution 10, 345–353. (doi:10.1016/j.tree.2010.01.007)

21. Kerr JT et al. 2015 Climate change impacts on bumblebees converge across continents. Science 349, 177–180. (doi:10.1126/science.aaa7031)

22. Soroye P, Newbold T, Kerr J. 2020 Climate change contributes to widespread declines among bumble bees across continents. Science 367, 685–688. (doi:10.1126/science.aax8591)

23. Powell E, Ratnayeke N, Moran NA. 2016 Strain diversity and host specificity in a specialized gut symbiont of honeybees and bumblebees. Molecular Ecology 25, 4461–4471. (doi:10.1111/mec.13787)

24. Zhang B, Leonard SP, Li Y, Moran NA. 2019 Obligate bacterial endosymbionts limit thermal tolerance of insect host species. Proceedings of the National Academy of Sciences 116, 24712–24718. (doi:10.1073/pnas.1915307116)

25. Heinrich B, Esch H. 1994 Thermoregulation in bees. American Scientist 82, 164–170.

26. Southwick EE, Heldmaier G. 1987 Temperature Control in Honey Bee Colonies. BioScience 37, 395–399. (doi:10.2307/1310562)

27. Heinrich B. 1979 Bumblebee Economics. Cambridge, Massachusetts: Harvard University Press.

28. Heinrich B. 1972 Energetics of temperature regulation and foraging in a bumblebee, *Bombus terricola* Kirby. Journal of Comparative Physiology 77, 49–64. (doi:10.1007/BF00696519)

29. Heinrich B. 1972 Patterns of endothermy in bumblebee queens, drones and workers. J. Comp. Physiol. 77, 65–79. (doi:10.1007/BF00696520)

30. Heinrich B. 1974 Thermoregulation in bumblebees: I. Brood incubation by *Bombus vosnesenskii* queens. J. Comp. Physiol. 88, 129–140. (doi:10.1007/BF00695404)

31. Tan K, Yang S, Wang Z-W, Radloff SE, Oldroyd BP. 2012 Differences in foraging and broodnest temperature in the honey bees *Apis cerana* and *A. mellifera*. Apidologie 43, 618–623. (doi:10.1007/s13592-012-0136-y)

32. Fahrenholz L, Lamprecht I, Schricker B. 1989 Thermal investigations of a honey bee colony: thermoregulation of the hive during summer and winter and heat production of members of different bee castes. J Comp Physiol B 159, 551–560. (doi:10.1007/BF00694379)

33. Kešnerová L, Emery O, Troilo M, Liberti J, Erkosar B, Engel P. 2020 Gut microbiota structure differs between honeybees in winter and summer. ISME J 14, 801–814. (doi:10.1038/s41396-019-0568-8)

34. Koch H, Abrol DP, Li J, Schmid-Hempel P. 2013 Diversity and evolutionary patterns of bacterial gut associates of corbiculate bees. Molecular Ecology 22, 2028–2044. (doi:10.1111/mec.12209)

35. Powell JE, Martinson VG, Urban-Mead K, Moran NA. 2014 Routes of Acquisition of the Gut Microbiota of the Honey Bee Apis mellifera. Applied and Environmental Microbiology 80, 7378–7387. (doi:10.1128/AEM.01861-14)

36. Koch H, Schmid-Hempel P. 2011 Socially transmitted gut microbiota protect bumble bees against an intestinal parasite. Proceedings of the National Academy of Sciences 108, 19288–19292. (doi:10.1073/pnas.1110474108)

37. Palmer-Young EC, Ngor L, Nevarez RB, Rothman JA, Raffel TR, McFrederick QS. 2019 Temperature dependence of parasitic infection and gut bacterial communities in bumble bees. Environ Microbiol 21, 4706–4723. (doi:10.1111/1462-2920.14805)

38. Martinson VG, Moy J, Moran NA. 2012 Establishment of characteristic gut bacteria during development of the honeybee worker. Applied and Environmental Microbiology 78, 2830–2840.

39. Michener CD. 1979 Biogeography of the Bees. Annals of the Missouri Botanical Garden 66, 277. (doi:10.2307/2398833)

40. Kwong WK, Engel P, Koch H, Moran NA. 2014 Genomics and host specialization of honey bee and bumble bee gut symbionts. Proceedings of the National Academy of Sciences 111, 11509–11514. (doi:10.1073/pnas.1405838111)

41. Vogt FD. 1986 Thermoregulation in Bumblebee Colonies. I. Thermoregulatory versus Brood-Maintenance Behaviors during Acute Changes in Ambient Temperature. Physiological Zoology 59, 55–59.

42. Gardner KE, Foster RL, O’Donnell S. 2007 Experimental analysis of worker division of labor in bumblebee nest thermoregulation (*Bombus huntii*, Hymenoptera: Apidae). Behav Ecol Sociobiol 61, 783–792. (doi:10.1007/s00265-006-0309-7)

43. Weidenmüller A, Kleineidam C, Tautz J. 2002 Collective control of nest climate parameters in bumblebee colonies. Animal Behaviour 63, 1065–1071. (doi:10.1006/anbe.2002.3020)

44. Jones JC, Myerscough MR, Graham S, Oldroyd BP. 2004 Honey bee nest thermoregulation: Diversity promotes stability. Science 305, 402–404. (doi:10.1126/science.1096340)

45. Steele MI, Kwong WK, Whiteley M, Moran NA. 2017 Diversification of Type VI Secretion System Toxins Reveals Ancient Antagonism among Bee Gut Microbes. mBio 8, e01630–17. (doi:10.1128/MBIO.01630-17)

46. Zheng H, Nishida A, Kwong WK, Koch H, Engel P, Steele MI, Moran NA. 2016 Metabolism of toxic sugars by strains of the bee gut symbiont *Gilliamella apicola*. mBio 7, e01326–16. (doi:10.1128/mBio.01326-16)

47. Williams P, Thorp R, Richardson L, Colla S. 2014 Bumble Bees of North America. Princeton, NJ: Princeton University Press.

48. R Core Team. 2016 R: A language and environment for statistical computing. Vienna, Austria: R Foundation for Statistical Computing. See http://www.r-project.org.

49. Sprouffske K, Wagner A. 2016 Growthcurver: an R package for obtaining interpretable metrics from microbial growth curves. BMC Bioinformatics 17, 172. (doi:10.1186/s12859-016-1016-7)

50. Näpflin K, Schmid-Hempel P. 2018 Host effects on microbiota community assembly. Journal of Animal Ecology 87, 331–340. (doi:10.1111/1365-2656.12768)

51. Sauers LA, Sadd BM. 2019 An interaction between host and microbe genotypes determines colonization success of a key bumble bee gut microbiota member. Evolution 73, 2333–2342. (doi:10.1111/evo.13853)

52. Klinger EG, Camp AA, Strange JP, Cox-Foster D, Lehmann DM. 2019 *Bombus* (Hymenoptera: Apidae) Microcolonies as a Tool for Biological Understanding and Pesticide Risk Assessment. Environmental Entomology 48, 1249–1259.

53. Leza M, Watrous KM, Bratu J, Woodard SH. 2018 Effects of neonicotinoid insecticide exposure and monofloral diet on nest-founding bumblebee queens. Proceedings of the Royal Society B: Biological Sciences 285, 20180761. (doi:10.1098/rspb.2018.0761)

54. Pinheiro J, Bates D, DebRoy S, Sarkar D, R Core Team. 2019 Linear and Nonlinear Mixed Effects Models. See https://CRAN.R-project.org/package=nlme.

55. Palmer-Young EC, Raffel TR, McFrederick QS. In press. Temperature-mediated inhibition of a bumblebee parasite by an intestinal symbiont. Proceedings of the Royal Society B 285, 2018 2041. (doi:http://dx.doi.org/10.1098/rspb.2018.2041)

56. Jones JC. 2004 Honey Bee Nest Thermoregulation: Diversity Promotes Stability. Science 305, 402–404. (doi:10.1126/science.1096340)

57. Heinrich B. 1976 Heat exchange in relation to blood flow between thorax and abdomen in bumblebees. The Journal of experimental biology 64, 561–85.

58. Hamblin AL, Youngsteadt E, López-Uribe MM, Frank SD. 2017 Physiological thermal limits predict differential responses of bees to urban heat-island effects. Biology Letters 13, 20170125. (doi:10.1098/rsbl.2017.0125)

59. Oyen KJ, Giri S, Dillon ME. 2016 Altitudinal variation in bumble bee (*Bombus*) critical thermal limits. Journal of Thermal Biology 59, 52–57. (doi:10.1016/j.jtherbio.2016.04.015)

60. Kovac H, Käfer H, Stabentheiner A, Costa C. 2014 Metabolism and upper thermal limits of *Apis mellifera carnica* and *A. m. ligustica*. Apidologie 45, 664–677. (doi:10.1007/s13592-014-0284-3)

61. Käfer H, Kovac H, Stabentheiner A. 2012 Upper thermal limits of honeybee *(Apis mellifera)* and yellowjacket *(Vespula vulgaris)* foragers. Mitt. Dtsch. Ges. Allg. Angew. Ent. 18, 267–270.

62. Ken T, Hepburn HR, Radloff SE, Yusheng Y, Yiqiu L, Danyin Z, Neumann P. 2005 Heat-balling wasps by honeybees. Naturwissenschaften 92, 492–495. (doi:10.1007/s00114-005-0026-5)

63. Miller-Struttmann NE et al. 2015 Functional mismatch in a bumble bee pollination mutualism under climate change. Science 349, 1541–1544. (doi:10.1126/science.aab0868)

64. Rasmont P, Iserbyt S. 2012 The Bumblebees Scarcity Syndrome: Are heat waves leading to local extinctions of bumblebees (Hymenoptera: Apidae: Bombus)? Annales de la Société entomologique de France (N.S.) 48, 275–280. (doi:10.1080/00379271.2012.10697776)

65. Woodard SH. 2017 Bumble bee ecophysiology: integrating the changing environment and the organism. Current Opinion in Insect Science 22, 101–108. (doi:10.1016/j.cois.2017.06.001)

66. Anbutsu H et al. 2017 Small genome symbiont underlies cuticle hardness in beetles. Proceedings of the National Academy of Sciences 114, E8382–E8391.

67. Engel P, James RR, Koga R, Kwong WK, McFrederick QS, Moran NA. 2013 Standard methods for research on *Apis mellifera* gut symbionts. Journal of Apicultural Research 52, 1–24. (doi:10.3896/IBRA.1.52.4.07)

68. Wertz JT, Breznak JA. 2007 *Stenoxybacter acetivorans* gen. nov., sp. nov., an Acetate-Oxidizing Obligate Microaerophile among Diverse O2-Consuming Bacteria from Termite Guts. Appl. Environ. Microbiol. 73, 6819–6828. (doi:10.1128/AEM.00786-07)

69. Benjamin P. Oldroyd, Siriwat Wongsiri. 2006 Asian Honey Bees: Biology, Conservation, and Human Interactions. Cambridge, Massachusetts: Harvard University Press.

70. D’Angelo C, Hume BCC, Burt J, Smith EG, Achterberg EP, Wiedenmann J. 2015 Local adaptation constrains the distribution potential of heat-tolerant *Symbiodinium* from the Persian/Arabian Gulf. ISME J 9, 2551–2560. (doi:10.1038/ismej.2015.80)

71. Mueller UG et al. 2011 Evolution of cold-tolerant fungal symbionts permits winter fungiculture by leafcutter ants at the northern frontier of a tropical ant-fungus symbiosis. Proceedings of the National Academy of Sciences 108, 4053–4056. (doi:10.1073/pnas.1015806108)

72. Okada S, Gordon DM. 2001 Host and geographical factors influence the thermal niche of enteric bacteria isolated from native Australian mammals. Molecular Ecology 10, 2499–2513. (doi:10.1046/j.0962-1083.2001.01384.x)

73. Cameron SA. 1985 Brood care by male bumble bees. Proceedings of the National Academy of Sciences 82, 6371–6373. (doi:10.1073/pnas.82.19.6371)

74. Heinrich B. 1972 Physiology of Brood Incubation in the Bumblebee Queen, *Bombus vosnesenskii*. Nature 239, 223–225.

75. Kelemen E, Dornhaus A. 2018 Lower temperatures decrease worker size variation but do not affect fine-grained thermoregulation in bumble bees. Behav Ecol Sociobiol 72, 170. (doi:10.1007/s00265-018-2577-4)

76. O’Donnell S, Foster RL. 2001 Thresholds of Response in Nest Thermoregulation by Worker Bumble Bees, *Bombus bifarius nearcticus* (Hymenoptera: Apidae). Ethology 107, 387–399. (doi:10.1046/j.1439-0310.2001.00668.x)

77. Livesey JS et al. 2019 The power and efficiency of brood incubation in queenless microcolonies of bumble bees (*Bombus terrestris* L.). Ecol Entomol 44, 601–609. (doi:10.1111/een.12736)

78. Burke G, Fiehn O, Moran N. 2010 Effects of facultative symbionts and heat stress on the metabolome of pea aphids. ISME Journal 4, 242–252. (doi:10.1038/ismej.2009.114)

79. Ma L, Calfee BC, Morris JJ, Johnson ZI, Zinser ER. 2018 Degradation of hydrogen peroxide at the ocean’s surface: the influence of the microbial community on the realized thermal niche of *Prochlorococcus*. ISME J 12, 473–484. (doi:10.1038/ismej.2017.182)

80. Bordenstein SR, Bordenstein SR. 2011 Temperature Affects the Tripartite Interactions between Bacteriophage WO, *Wolbachia*, and Cytoplasmic Incompatibility. PLoS ONE 6, e29106. (doi:10.1371/journal.pone.0029106)

81. Marquez LM, Redman RS, Rodriguez RJ, Roossinck MJ. 2007 A Virus in a Fungus in a Plant: Three-Way Symbiosis Required for Thermal Tolerance. Science 315, 513–515. (doi:10.1126/science.1136237)

82. Deboutte W, Beller L, Yinda CK, Maes P, de Graaf DC, Matthijnssens J. 2020 Honey-bee–associated prokaryotic viral communities reveal wide viral diversity and a profound metabolic coding potential. Proc Natl Acad Sci USA 117, 10511–10519. (doi:10.1073/pnas.1921859117)

83. Bonilla-Rosso G, Steiner T, Wichmann F, Bexkens E, Engel P. 2020 Honey bees harbor a diverse gut virome engaging in nested strain-level interactions with the microbiota. Proc Natl Acad Sci USA 117, 7355–7362. (doi:10.1073/pnas.2000228117)

84. Crall JD et al. 2018 Neonicotinoid exposure disrupts bumblebee nest behavior, social networks, and thermoregulation. Science 362, 683–686. (doi:10.1126/science.aat1598)

85. Lemoine MM, Engl T, Kaltenpoth M. 2020 Microbial symbionts expanding or constraining abiotic niche space in insects. Current Opinion in Insect Science 39, 14–20. (doi:10.1016/j.cois.2020.01.003)

86. Lewis Z, Lizé A. 2015 Insect behaviour and the microbiome. Current Opinion in Insect Science 9, 86–90. (doi:10.1016/j.cois.2015.03.003)

87. Sherwin E, Bordenstein SR, Quinn JL, Dinan TG, Cryan JF. 2019 Microbiota and the social brain. Science 366, eaar2016. (doi:10.1126/science.aar2016)

